# Time dependent proinflammatory responses shape virus interference during coinfections of influenza A virus and influenza D virus

**DOI:** 10.1101/2021.03.14.435363

**Authors:** Minhui Guan, Sherry Blackmon, Alicia K. Olivier, Xiaojian Zhang, Liyuan Liu, Amelia Woolums, Mark A. Crenshaw, Shengfa F. Liao, Richard Webby, William Epperson, Xiu-Feng Wan

**Author notes:** Corresponding author: Dr. Xiu-Feng Wan by. Contributed equally to this study (Minhui Guan, Sherry Blackmon). Author order was determined by a mutual agreement and in order of increasing seniority.

## Abstract

Both influenza A virus (IAV) and influenza D virus (IDV) are enzootic in pigs. IAV causes approximately 100% morbidity with low mortality, whereas IDV leads to only mild respiratory diseases in pigs. In this study, we performed a series of coinfection experiments *in vitro* and *in vivo* to understand how IAV and IDV interact and cause pathogenesis during coinfection. Results showed that IAV inhibited IDV replication when infecting swine tracheal epithelial cells (STEC) with IAV 24- or 48-hours prior to IDV inoculation, and that IDV suppressed IAV replication when IDV preceded IAV inoculation by 48 hours. Virus interference was not identified during simultaneous IAV/IDV infections or with 6 hours between the two viral infections, regardless of their order. The interference pattern at 24- and 48-hours correlated with proinflammatory responses induced by the first infection, which was about 24-hours slower for IDV than IAV. The viruses did not interfere with each other if both infected the cells before proinflammatory responses were induced. Coinfection in pigs further demonstrated that IAV interfered both viral shedding and virus replication of IDV, especially in the upper respiratory tract. Clinically, coinfection of IDV and IAV did not show significant enhancement of disease pathogenesis, compared with the pigs infected with IAV alone. In summary, this study suggests that interference during coinfection of IAV and IDV is primarily due to the proinflammatory response and is therefore dependent on the time between infection, and the order of infection.

**Importance:** Both IAV and IDV are enzootic in pigs, and feral pigs have a higher risk for both IAV and IDV exposures than IDV exposure alone. This study suggests that in coinfection with IAV and IDV either virus can interfere with the replication of the other virus by stimulating proinflammatory responses; however, the proinflammatory response was 24 hours slower for IDV than IAV. *In vitro* there was no interference during simultaneous coinfection, regardless of infection order. Coinfection of IDV and IAV in pigs did not show enhanced pathogenesis, compared with those infected only with IAV. This study can facilitate our understanding of virus epidemiology and pathogenesis associated with IAV and IDV coinfection.

## Introduction

Influenza viruses are classified into types A, B, C and D according to genetic and antigenic properties of the nucleoprotein (NP) and matrix 1 (M1) genes (1, 2). Whereas influenza B and C viruses are documented to infect only humans and swine, influenza A virus (IAV) and influenza D virus (IDV) can infect a wide range of hosts. In addition to humans, IAV can infect pigs, horses, dogs, marine mammals (e.g. seals and whales), and a spectrum of avian species, including both wild birds and domestic poultry; IDV can infect domestic and feral swine, cattle, goats, sheep, camelids, buffalo, and equids (2–8), and a low level of human exposure to IDV is also documented (9). The influenza genome consists of negative-sense, single-stranded, segmented RNA. The genome of IAV contains eight segments, encoding at least 10 or even 14 proteins (10), whereas those of IDV contain seven segments, encoding at least nine proteins. Based upon the surface glycoproteins, hemagglutinin (HA) and neuraminidase (NA), IAV is further classified into 18 HA and 11 NA types (11, 12). Different from IAV, IDV encodes a hemagglutinin-esterase-fusion (HEF) surface glycoprotein that catalyzes receptor binding, cleavage, and membrane fusion (3, 13, 14), resembling the functions of HA and NA for IAV.

IAV can stimulate both innate and adaptive immune responses with variations that are host dependent. IAV induces the host innate immune response and promotes disease pathogenesis through non-structural (NS1) protein to inhibit TRIM25 ubiquitination, which is required for the activation of retinoic acid-inducible gene I (RIG-I) mediated interferon production (15). The RIG-I pathway, essential in epithelial cell interferon induction, is induced by preferentially binding viral RNA with a greater affinity for single-stranded RNA without 5’OH or 5’-methylguanosine cap (16). Presence of RIG-I in ducks (but not in chickens) induces proinflammatory responses and ultimately facilitates viral clearance in ducks during IAV infection (17). In humans, modulating the innate immune response via interferon inhibition often enhances virus production. However, in some cases the interferon response is too robust which may lead to a “cytokine storm” characterized by overproduction of interferon leading to upregulation of additional proinflammatory cytokines, excessive infiltration of the tissue by immune cells leading to tissue destruction. The cytokine storm was reported for the 1918 pandemic H1N1 virus (18) and H5N1 highly pathogenic avian influenza virus (19–22).

IDV infection in IDV seronegative calves cause mild pathogenicity in cattle (23), with mild respiratory disease with respiratory tract inflammation characterized by multifocal mild tracheal epithelial attenuation and neutrophil infiltration. In field studies, IDV has been associated with bovine respiratory disease (BRD) complex, a disease of significant economic burden. A previous study had no evidence of cattle coinfected with IDV and *Mannheimia haemolytica,* a pathogen commonly detected in BRD, to have worse clinical scores or lung pathology than animals infected with only *Mannheimia haemolytica* (24). In animal models, IDV replicated in the upper and lower respiratory tracts of pigs (3, 25), guinea pigs (26) and mice (27), and the overall clinical diseases in these animal models caused by a single IDV infection were mild. Nevertheless, the overall role of IDV in pathogenesis, especially during coinfections with other pathogens, including IAV, is still unclear.

Both IAV and IDV are documented to be enzootic in pigs, based on serological evidence from a set of feral swine sera samples collected in the U.S. from 2010–2013 where approximately 43%, were seropositive for IDV and IAV, suggesting the host-pathogen ecology may include coinfections (25). In addition, the seroprevalence rate of IDV in IAV-seropositive feral swine was more than twice that observed among IAV-negative feral swine, suggesting the possibility of virus interactions during IAV and IDV coinfection (25). The objectives of this study were to evaluate the interactions between IAV and IDV during coinfection and to evaluate the pathogenesis during IAV-IDV coinfection in influenza seronegative pigs. By using an *in vitro* system, we compared proinflammatory responses of IAV and IDV and further correlated these responses with the virus interference patterns and with the factors of infection order and infection time gap. Additionally, we evaluated clinical pathogenesis of IAV and IDV coinfection using a pig model.

## Results

### Both IAV and IDV stimulate proinflammatory responses but in a different speed

To compare proinflammatory responses stimulated by IAV and IDV, we evaluated both gene and protein expression in swine tracheal epithelial primary cells (STEC), which was kindly provided by Dr. Stacey Schultz-Cherry, for a set of proinflammatory markers, including type I interferon (IFN-β), type II interferon (IFN-γ), tumor necrosis factor alpha (TNF-α), DDX58 (retinoic acid-inducible gene I [RIG-I]), interleukin (IL)-1β, IL-4, IL-6, IL8, IL-10, IP-10 (also called CXCL10, interferon-γ-inducible protein 10, previously called IP-10), C–C chemokine ligand 5 (CCL5), and C-X-C Motif Chemokine Ligand 9 (CXCL9)] (Table 1), which were reported in IAV and/or IDV infection (28, 29). In all *in vitro* experiments, A/swine/Texas/A01104013/2012(H3N2) (sH3N2) and D/bovine/Mississippi/C00046N/2014 (D/46N) were used, and multiplicity of infection (MOI) of 0.001 and 0.1 were implemented for IAV and IDV, respectively.

**Table 1.**
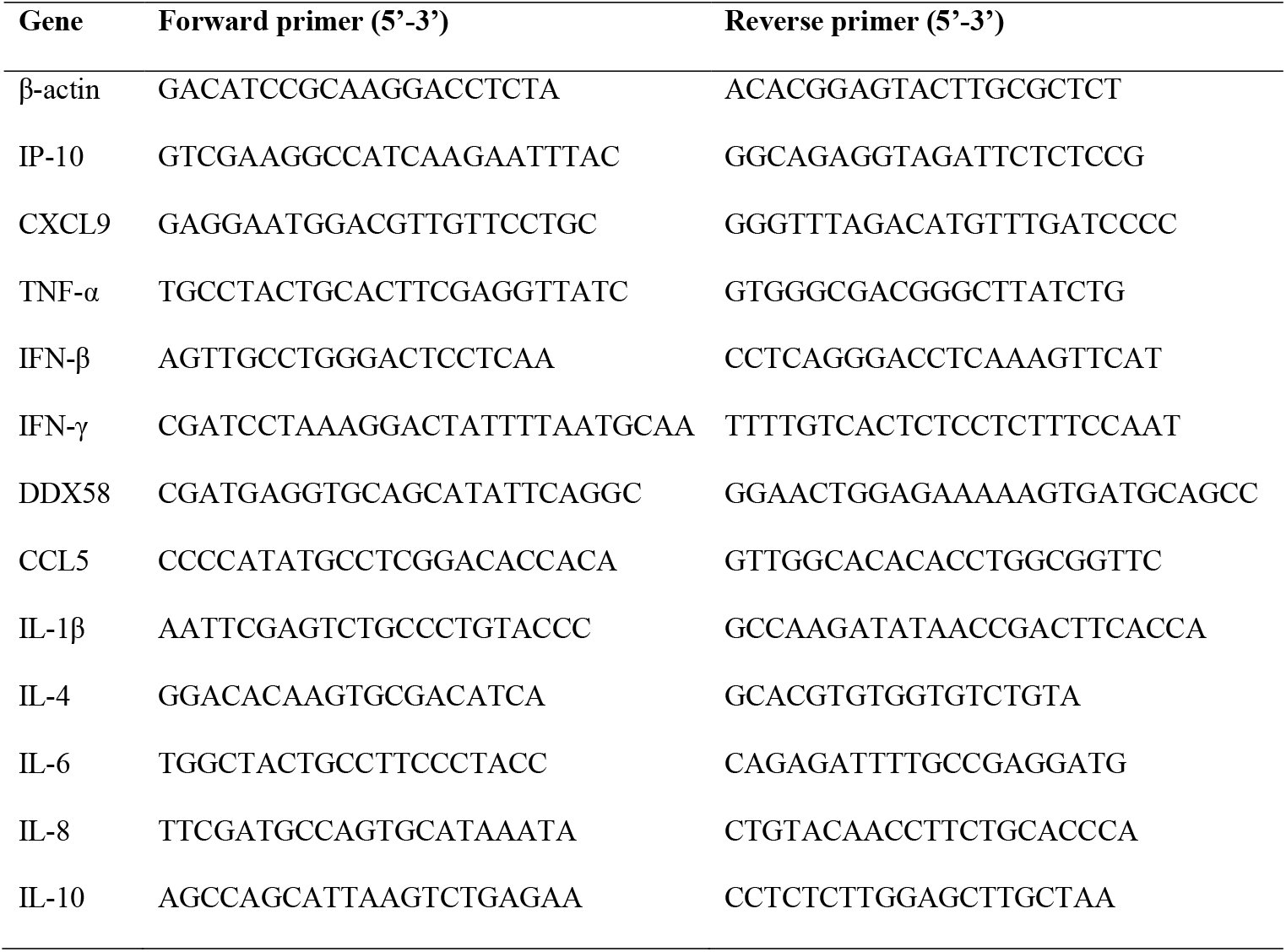
Primers used to quantify mRNA expression of proinflammatory markers in STEC.

For IAV infection, results from quantitative RT-PCR (qRT-PCR) showed that, compared with those in the negative control, the IP-10 and CXCL9 had the fastest and highest responses, with 9.92 (± 0.03; standard deviation)- and 5.89 (± 0.87) -fold increases at 24 hours post inoculation (hpi) and with 516.78 (± 110.48)- and 768.59 (± 330.43)-fold increases at 48 hpi, respectively. Gene expression of TNF-α, CCL5 and DDX58 were significantly increased at 48 hpi (12.86 ± 1.57; 20.18 ± 2.19; 28.84 ± 3.33, respectively) and remained elevated at 72 hpi whereas IFN-β and IL-6 increased at 48 hpi (15.49 ± 4.21; 2.87 ± 1.19, respectively) but rapidly decreased at 72 hpi. Gene expression of IFN-γ, IL-4, IL-8, IL-10, and IL-1β were not significantly changed (Fig 1).

**Fig 1.**
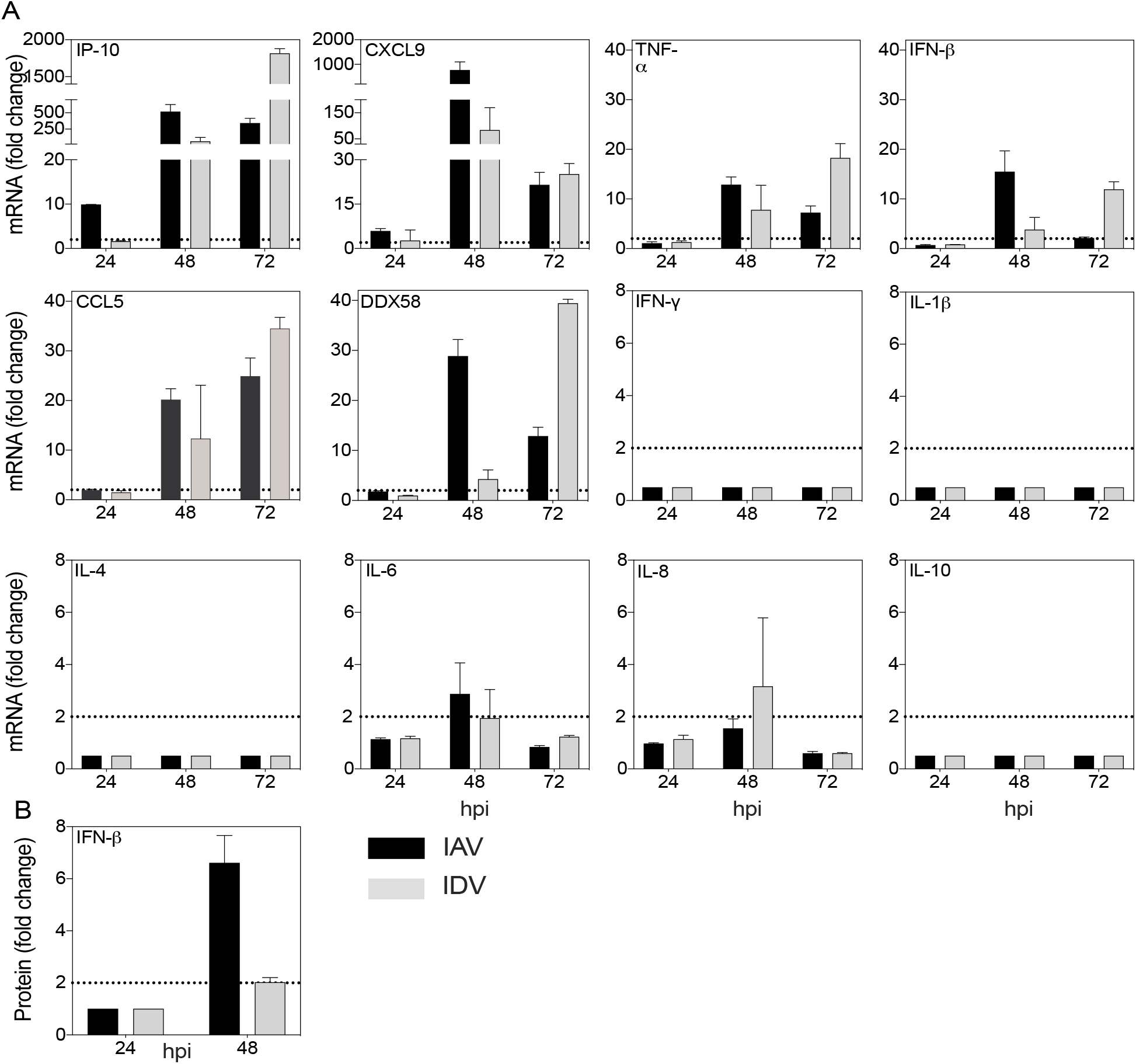
Proinflammatory responses stimulated by IAV and IDV at swine tracheal epithelial primary cells (STEC). Proinflammatory responses induced by IAV or IDV alone were showed in black and grey bars, respectively. A) The relative mRNA expressions were quantified by qPCR, and normalized by β-actin, a housekeeping gene. ΔΔCt (Ct is the cycle threshold) = ΔCt (sample) ([Ct _gene of interest_-Ct _housekeeping gene_] of infected cells) - ΔCt (negative) ([Ct_gene of interest_ -Ct _housekeeping gene_] of uninfected cells). The mean values of fold change (2^-ΔΔCt^) of triplicates and standard deviation were represented. B) The protein expression of IFN-β were quantified by ELISA assays. The fold change values of IFN-β’s protein level was calculated (infected samples/negative samples) and plotted as y axis. hpi, hours post inoculation.

For IDV infection, none of the proinflammatory markers we evaluated showed upregulated gene expression at 24 hpi. Similar to the IAV infection, the expression of six genes significantly started to increase at 48 hpi (IP-10, 62.20 ± 60.88; CXCL9, 83.68 ± 85.79; TNF-α, 7.78 ± 4.99; IFN-β, 3.79 ± 2.50; CCL5, 12.32 ± 10.76; DDX58, 4.24 ± 1.86) and remained elevated at 72 hpi (Fig 1). Among them, IP-10 and CXCL9 has the highest upregulated expression. Gene expression of IFN-γ, IL-4, IL-6, IL-8, IL-10, and IL-1β was not significantly affected.

To further validate the proinflammatory responses at the protein level, we quantified IFN-β in cell supernatants harvested at 24 and 48 hpi by ELISA assay. Results showed that IFN-β was upregulated with 6.61 (±1.05)- and 2.02 (± 0.18)- folds for IAV and IDV infections at 48 hpi, respectively, both of which correlate to increased mRNA expression.

In summary, both IAV and IDV stimulated similar proinflammatory responses, including IP-10, CXCL9, TNF-α, CCL5, DDX58, CXCL9, and IFN-β with a similar level of upregulation; however, the proinflammatory responses by IAV appeared approximately 24 hours earlier than IDV.

### IAV and IDV interfere the replication of each other during coinfection

We hypothesize that proinflammatory responses stimulated by IAV and IDV interfere with virus replication during coinfection, and thus interference will be dependent on the order and time gap of virus infection. To test this hypothesis, we performed a series of coinfection experiments in STECs by 1) simultaneous inoculation of IAV and IDV (A+D); 2) sequential inoculations with IAV followed by IDV (A-D groups) with time gaps of 6 (A-D-6h), 24 (A-D-24h) and 48 hours (A-D-48h), and 3) sequential inoculations with IDV followed by IAV (D-A groups) with time gaps of 6 hours (D-A-6h), 24 (D-A-24h) and 48 hours (D-A-48h). The infection groups of IAV (A-mock) or IDV (D-mock) alone were included as mock controls. The viral copies of IAV and IDV were quantified using IAV and IDV matrix-gene (M) specific qRT-PCR.

IAV reached titers of 2.99 (± 0.38), 5.00 (± 0.26), and 5.67 (± 0.27) log_10_ copies/μl at 24, 48, and 72 hpi in A-mock, respectively; correspondingly, IDV had 4.68 (± 0.26), 5.11 (± 0.22), and 5.19 (± 0.21) log_10_ copies/μl in D-mock. In the A+D group, IAV and IDV reached the growth plateau with a titer of 5.01 (± 0.27) and 5.03 (± 0.22) log_10_ copies/μl at 48 hpi, respectively. Both the titers of either IAV or IDV in A+D were not statistically significant different from those corresponding titers in A-mock (*p* = 0.842) or D-mock (*p* >0.9999) (Fig 2A).

**Fig 2.**
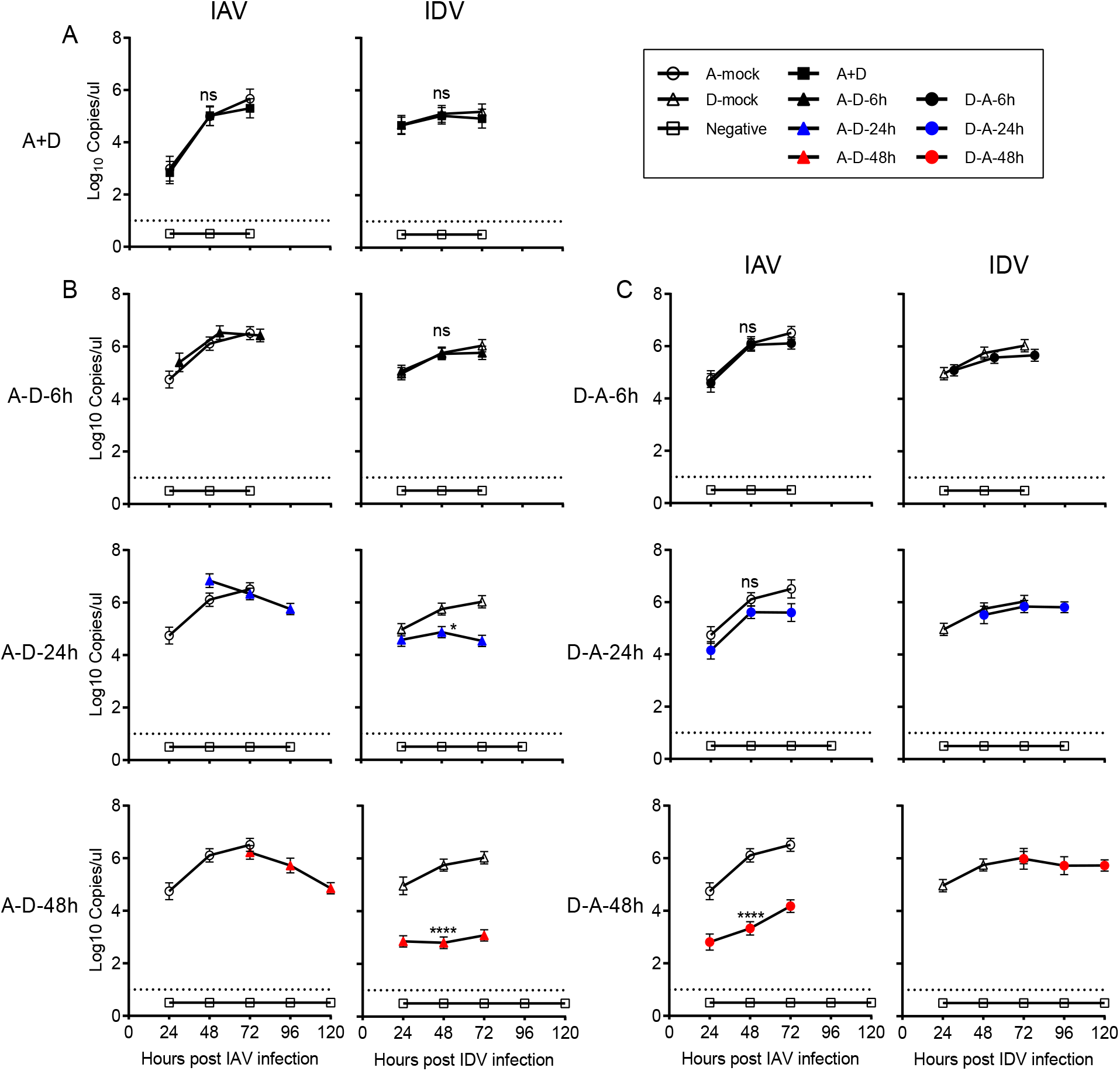
Growth kinetics of coinfecting IAV and IDV in STEC. A) Simultaneous coinfections (A+D); B) sequential infections of IAV infection followed by IDV (A-D) with a time gap of 6 (A-D-6h), 24 (A-D-24h) or 48 (A-D-48h) hours; C) sequential infections of IDV infection followed by IAV with a time gap of 6 (D-A-6h), 24 (D-A-24h) or 48 (D-A-48h) hours. Growth kinetics assays were performed in STEC at 37°C in triplicates with a MOI of 0.001 of IAV and 0.1 of IDV, respectively. At each time point, supernatant was collected and replaced, and RNA copy numbers was determined by qRT-PCR. The left panel of each subfigure showed the detection of IAV whereas the right panel showed the detection of IDV. The x axis of left panel in each subfigure represents hours post IAV infection whereas the right panel represents hours post IDV infection of the corresponding samples. The mean copy numbers and standard deviation were calculated for each experimental replicate at each time, and the dotted line denoted the detecting limit of 1. One-way repeated measures ANOVA were performed to compare between the single infection and coinfection groups, and significant differences were presented (\**p*≤0.05, \*\**p*<0.0021, \*\*\**p*<0.0002, \*\*\*\**p*<0.0001) and not significant (*p*>0.05) differences as ns.

For the A-D sequential infection groups, at 48 hpi, IDV had 5.72 (± 0.22), 4.87 (± 0.21), and 2.80 (± 0.22) log_10_ copies/ul in A-D-6h, A-D-24h, and A-D-48h, respectively. Compared with those at the A-mock group, there were a 1.08 (± 0.12)-fold change at A-D-6h, a 7.61 (±0.29)-fold decrease at A-D-24h, and a 882.42 (±16.61)-fold decrease at A-D-48h. Statistical analyses showed that the IDV titers were significantly lower in A-D-24h (*p* = 0.0204) and A-D-48h (*p* < 0.0001) than in D-mock, but no significant difference was identified between A-D-6h and D-mock (*p* > 0.9999) (Fig 2B). The titers of IAV in all three A-D groups were not statistically different from those of A-mock.

For the D-A sequential infection groups, at 48 hpi, the IAV had 6.05 (± 0.24), 5.62 (± 0.23), and 3.34 (± 0.25) log_10_ copies/ul in D-A-6h, D-A-24h, and D-A-48h, respectively. Compared with those at the A-mock group, there was a 590.44 (± 12.39)-fold decrease in the viral titers of IAV at 48 hpi in the D-A-48h groups. Statistical analyses showed that the IAV titers were significantly lower in D-A-48h (*p* < 0.0001) but not affected in D-A-6h (*p* >0.9999) or D-A-24h (*p* = 0.056) (Fig 2C). The titers of IDV in the A-D groups were not statistical different from those from the D-mock group.

Taken together, our results suggest that IAV inhibited IDV replication in STEC when IAV preceded IDV inoculation by 24 or more hours. IDV inhibited IAV replication when IDV preceded IDA inoculation by 48 hours. The viral interference correlated with the speed of the proinflammatory responses induced by the first infection, which was about 24-hours slower for IDV than IAV, and the viruses did not interfere when cells were coinfected simultaneously, before proinflammatory responses were induced, validating our hypotheses.

### Coinfection of IAV and IDV in pigs limited replication of IDV but not IAV in upper respiratory tracts

To evaluate the pathogenesis during coinfection of IAV and IDV, we simultaneously intranasally inoculated pigs with 10^6^ TCID_50_/ml of sH3N2 and the same amount of D/46N, both of which resulted in effective virus replication and shedding in pigs (25, 30). Single infection groups (A-mock [n = 5] or D-mock [n = 7]) and a control group inoculated with sterile PBS (Negative group; n = 6) were included as controls.

Results showed in the single infections, 5/5 pigs shed IAV and 6/7 pigs shed IDV at 3, 4 and/or 5 dpi, respectively. In A-mock, viral shedding peaked at 3 dpi (6.10 log_10_ copies/ml); whereas, virus shedding in D-mock peaked at 5 dpi (4.72 log_10_ copies/ml). In the coinfection group A+D, 7/7 shed IAV which peaked at 3 dpi (5.33 log_10_ copies/ml), but only 1/7 pigs shed IDV, which peaked at 4 dpi with low viral copies (3.95 log_10_ copies/ml). To evaluate how coinfection affects viral shedding, we compared viral shedding between A-mock and A+D and found that the IAV shedding showed no significant difference during the five days (*p* = 0.1262). Of interest, pigs in the coinfection A+D group showed significantly decreased shedding of IDV, compared to D-mock (*p* < 0.0001) (Fig 3).

**Fig 3.**
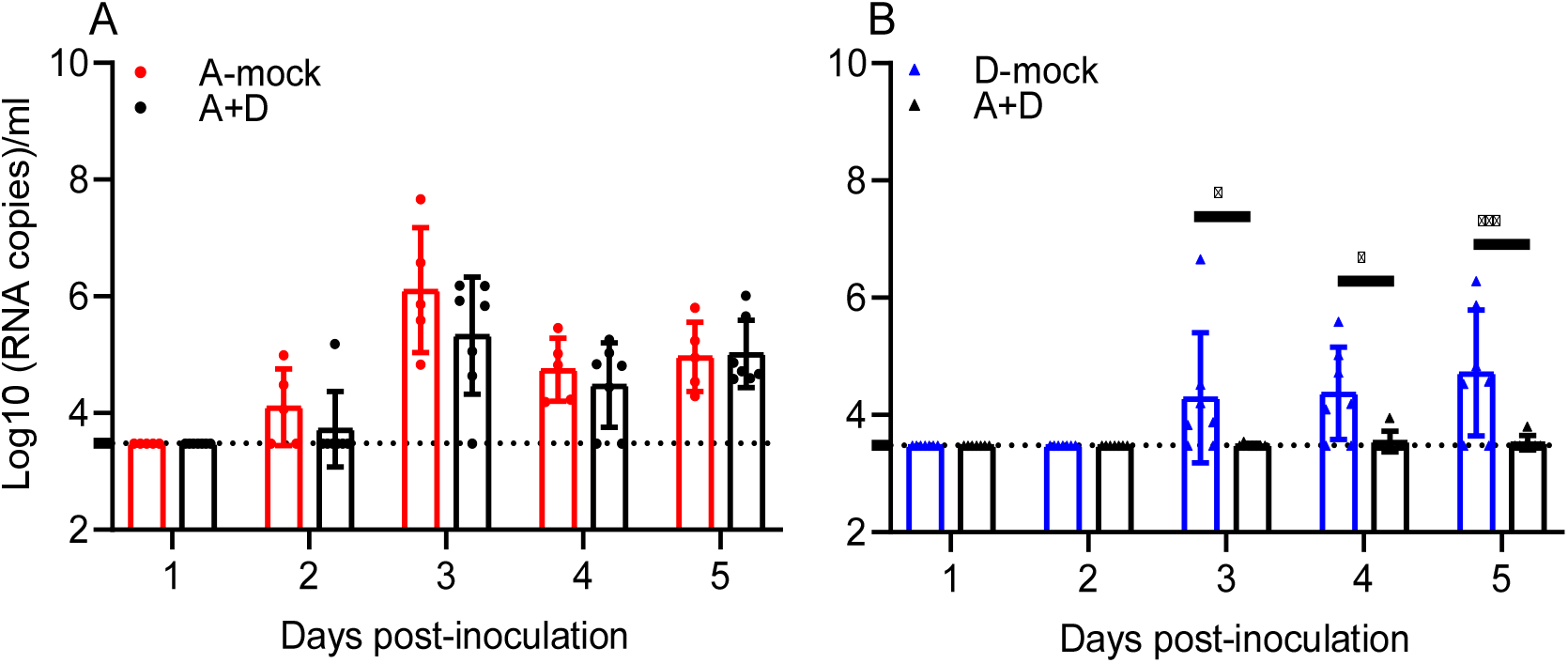
Viral shedding in the pigs from the IAV and IDV coinfection experiment. A) Viral titers of IAV; B) Viral titers of IDV. Viral loads in each nasal wash were quantified by qRT-PCR and represented as log_10_ (RNA copies)/ml. Each bar represents the mean values per group and standard deviation. Each data point indicated one sample. The dashed line indicates the limit of detection of 3.48 log_10_ copies/ml. Samples from different treatment groups are differentiated in colors: A-mock group in red, D-mock group in blue and A+D group in black. No IAV was detected in the negative group and D-mock group. No IDV was detected in the negative group and A-mock group (not shown). Two-way ANOVA analysis was performed to compare between the single infection and coinfection groups, and significant differences were presented (\**p*≤0.05, \*\**p*<0.0021, \*\*\**p*<0.0002, \*\*\*\**p*<0.0001) and not significant (*p*>0.05) differences as ns.

To further evaluate the coinfection on tissue dependent viral replication, all pigs were euthanized on 5 dpi and the viral load in the tissues of the respiratory tree were quantified. The fourteen respiratory tract tissues were categorized into four groups: 1) turbinate [rostral (RT), middle (MT), ethmoid turbinate (ET)], 2) trachea [upper (TR-U), middle (TR-M), distal (TR-D)], 3) soft palate (SP), and 4) lower respiratory tract [bronchus (BR), lung left cranial (LCR) and caudal (LCD), and right cranial (RCR), caudal (RCD), middle (RM) and accessory (RA) lobes]. In tissues from both A-mock and A+D, all pigs were positive for IAV with a detection limit of 4.03 log_10_ copies/g. There was no significant difference between A-mock and A+D for respiratory tissue IAV replication (*p* = 0.8214) (Fig 4A).

**Fig 4.**
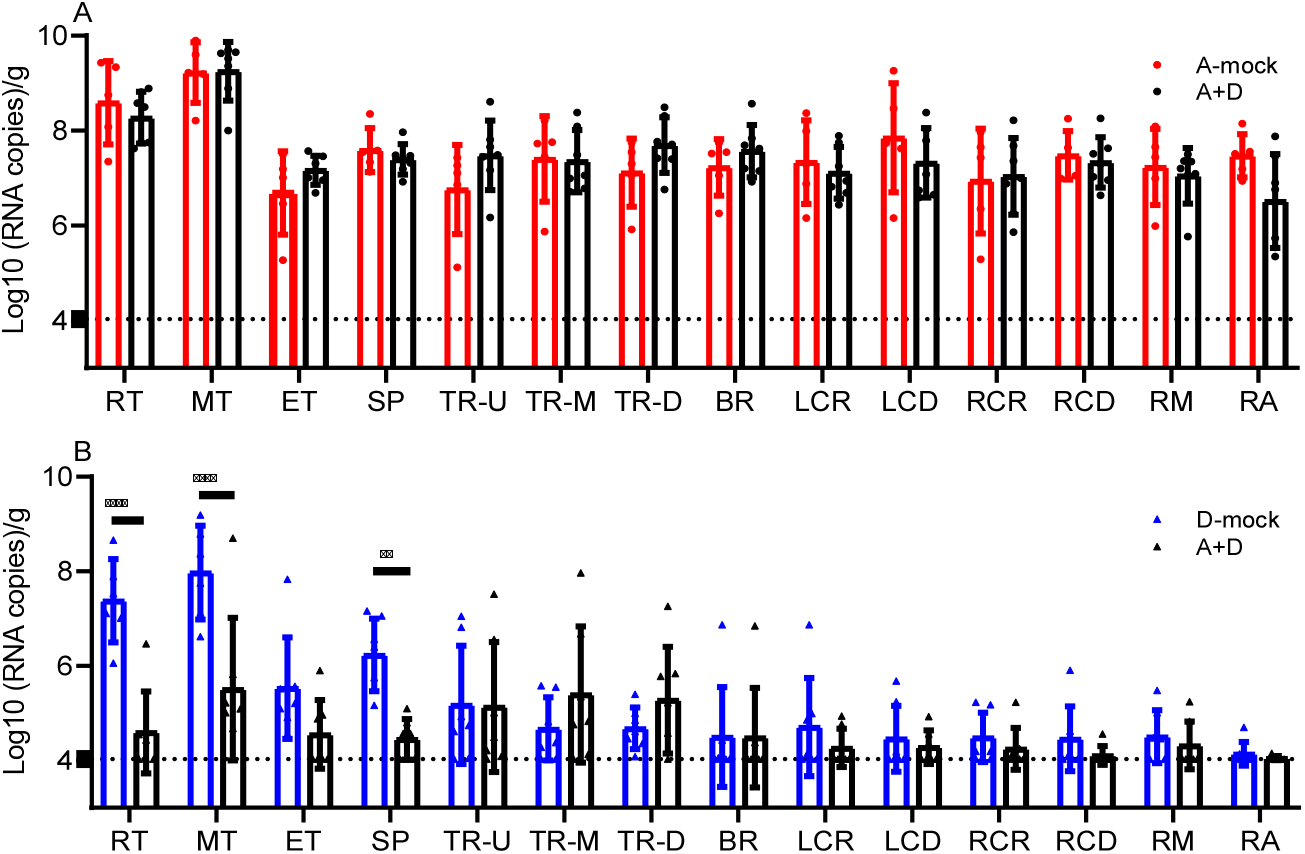
Viral titers in the respiratory tract tissues of the pigs from the IAV and IDV coinfection experiment. A) Viral titers of IAV; B) Viral titers of IDV. Viral loads were quantified by qRT-PCR and represented as log_10_ (RNA copies)/g. Each bar represents the mean values per group and standard deviation. Each data point indicated one sample. The dashed line indicates the limit of detection of 4.03 log_10_ copies/g. Samples from different treatment groups are differentiated in colors: A-mock group in red, D-mock group in blue and A+D group in black. No IAV was detected in the negative group and D-mock group. No IDV was detected in the negative group and A-mock group (not shown). Two-way ANOVA analysis was performed to compare between the single infection and coinfection groups, and significant differences were presented (\**p*≤0.05, \*\**p*<0.0021, \*\*\**p*<0.0002, \*\*\*\**p*<0.0001) and not significant (*p*>0.05) differences as ns. Abbreviations: rostral turbinate (RT), middle turbinate (MT), ethmoid turbinate (ET), soft palate (SP), upper trachea (TR-U), middle trachea (TR-M), distal trachea (TR-D), bronchus (BR), left cranial lung (LCR), left caudal lung (LCD), right cranial lung (RCR), right caudal lung (RCD), right middle lung (RM) and right accessory lung (RA).

All pigs in the groups D-mock and A+D were positive for IDV. IDV copy number was lowest in the lower respiratory tract at 4.46 and 4.25 log_10_ copies/g, for single and coinfection groups, respectively. In the single infection IDV group, IDV copy number was the highest in the turbinate average at 6.96 log_10_ copies/g and less in the trachea at 4.84 log_10_copies/g (*p* < 0.001) and lower respiratory tract at 4.46 log_10_copies/g (*p* < 0.001). Differences were also significantly different between the soft palate (6.23 log_10_ copies/g) and lower respiratory tract (4.46 log_10_ copies/g) (*p* < 0.001) and trachea (4.84 log_10_ copies/g) (*p* = 0.001). In A+D, IDV copy number was higher in the trachea (5.27 log_10_ copies/g) than in the lower respiratory tract (4.25 log_10_ copies/g) (*p* = 0.032). There were no other differences among tissues in the coinfection group A+D (*p* > 0.137). IDV copy number was lower in the RT, MT, and SP in A+D than D-mock. Specifically, there were significant differences between the RTs with 4.59 vs 7.38 log_10_copies/g (*p* <0.0001), MTs with 5.51 vs 7.97 log_10_copies/g (*p* <0.0001) and the SPs with 4.45 vs 6.23 log_10_ copies/g (*p* = 0.0018) for A+D and D-mock, respectively. The trachea and lower respiratory tract showed no differences in IDV viral copy number when comparing D-mock and A+D (all *p* > 0.05) (Fig 4B).

Taken together, simultaneous co-inoculation of IAV and IDV in pigs significantly reduced viral shedding of IDV and viral replication of IDV in the tissues of the upper respiratory tract, but coinfection did not affect replication of IAV.

### Coinfection of IAV and IDV virus did not significantly enhance disease pathogenesis

Pigs in all virally inoculated groups had elevated body temperatures (A-mock, D-mock, and A+D) and lymphopenia. At 3 and 4 dpi, rectal temperatures were slightly higher in the single infection groups than the coinfection group; however, there was no statistical difference due to a large variation across pigs in the negative control group. Histologic evaluation of all tissues from the respiratory tract showed no significant differences between treatment groups. This is largely due to chronic inflammatory changes being present in the turbinate, trachea and lung. There was no significant acute inflammatory response in any of the tissues. Chronic tracheal inflammation was characterized by mucosal and submucosal infiltrates of lymphocytes, plasma cells and fewer eosinophils (Fig 5). In some sections of trachea apoptotic cells were frequent, but not significantly different between treatment groups (5D). In all lung sections, including control tissues, the interstitium was moderately thickened due to increased numbers of interstitial macrophages, eosinophils, and fewer lymphocytes. There was no evidence of bronchiolar epithelial loss or exudate within the lumen of the airways. The chronic inflammatory changes are thought to be due to the environmental housing and chronic antigenic stimulation.

**Fig 5.**
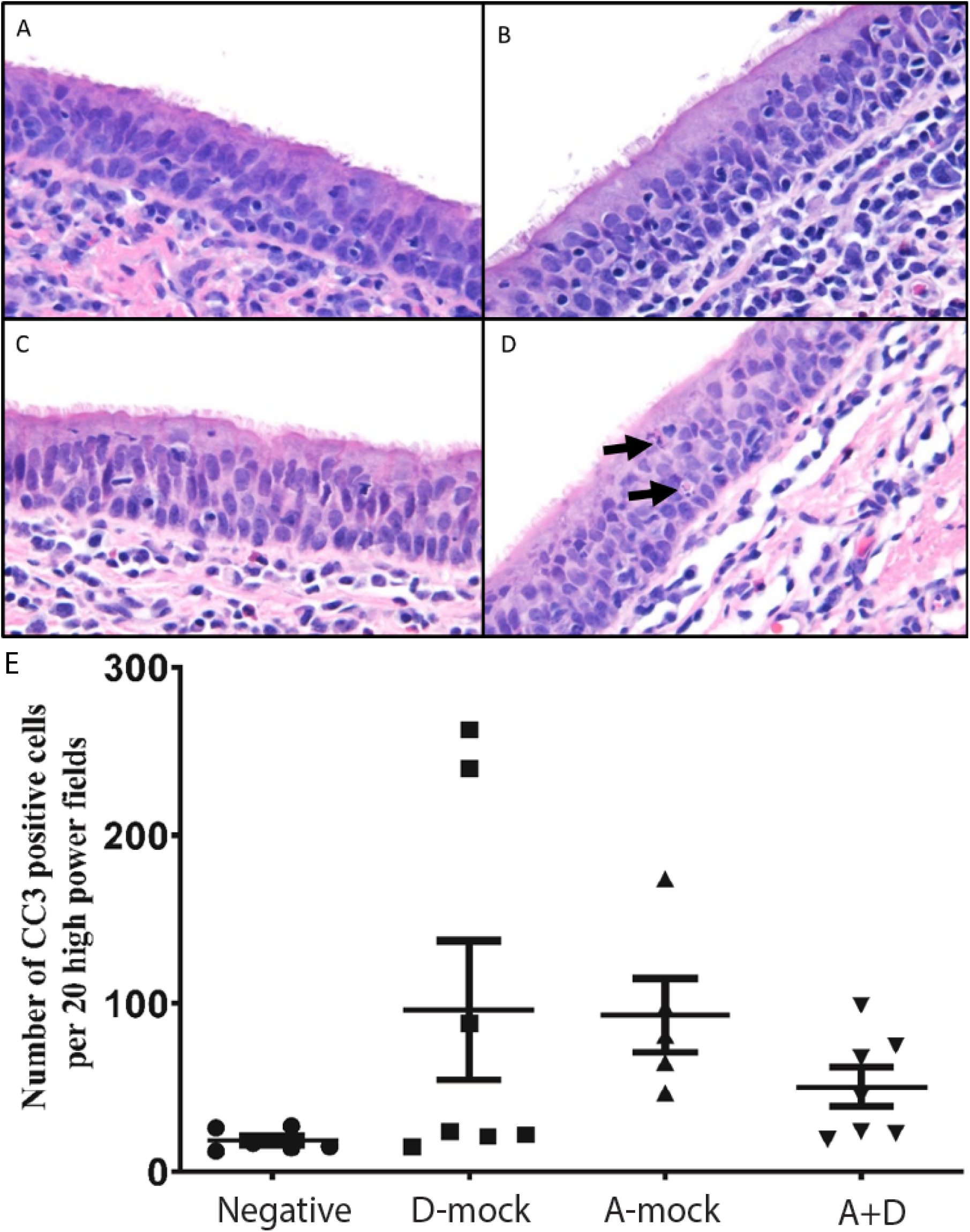
Hematoxylin and eosin stain staining of tracheas of pigs. A) Negative control, in which pigs were inoculated with sterile PBS; B) D-mock, in which pigs were inoculated with D/46N alone; C) A-mock, in which pigs were inoculated with sH3N2 alone; D) A+D, in which pigs were inoculated simultaneously with D/46N and sH3N2; and E) Cleaved caspase 3 staining in the trachea of pigs. All tracheal tissues showed chronic lymphoplasmacytic inflammation within the mucosa and submucosa. Apoptotic bodies (arrows) were frequently observed (5D, arrows), however no significantly different between treatment groups and varied between pigs (5E).

IAV and influenza B virus, like other viral pathogens, induce apoptosis *in vitro* and *in vivo*, and apoptosis is associated pathogenesis of influenza viruses (31, 32). Cleaved caspase-3 (CC3) is a downstream effector caspase and an important regulator of apoptosis, activation of Caspase 3 is shown to be essential for efficient influenza virus propagation (33). To further examine apoptosis associated pathogenesis among the virally infected groups, we determined examined and semi-quantified CC3 via immunohistochemistry in the tracheal tissues. In the A-mock group, the mean number of CC3 positive cells was 93 (+ 22 SEM; range: 47-174) which was higher than mean number of CC3 positive cells in the control group 19 (+ 3 SEM; range 12-27) (*p* ≤0.05) (Fig 5E). In the D-mock group, the mean number of CC3 positive cells was 96 (+ 41 SEM; range: 15-248). Of note, in the D-mock group, four pigs had low numbers of CC3 positive cells, but the counts were high for pigs 61 and 71. Interestingly, pig 71 was the only pig in the D-mock group that was positive for all respiratory tract tissues sampled. In the A+D group, the mean was 50 (+12 SEM; range: 19-99). Nevertheless, no statistical significances were found for the number of CC3 positive cells among A-mock, D-mock, and A+D groups.

Taken together, the coinfection did not lead to significant differences in the clinical signs or pathology compared to single infection.

## Discussion

The objectives of this study were to evaluate the interactions between IAV and IDV during coinfection, and to evaluate the pathogenesis during coinfection in influenza seronegative pigs. Our *in vitro* data showed that proinflammatory responses were stimulated by IDV at 48 hpi was 24 hours delayed compared to proinflammatory responses by IAV, although the expression levels and the associated genes were similar between IAV and IDV infections. Therefore, interference by IAV and IDV depends on both infection order and infection time gap, with no interference observed with simultaneous infection in STEC. In addition to STEC, experiments were also performed in MDCK cells, and the results were similar to those observed in STEC (data not shown). An animal challenge with IAV and IDV coinfection showed IDV nasal shedding and viral replication in nasal turbinate and soft palate were decreased during coinfection, suggesting simultaneous coinfection may have antagonistic effects on IDV viral replication *in vivo.* By examining viral shedding, we found that pigs infected with sH3N2 shed viruses at 2 dpi and peaked at 3 dpi whereas those infected D/46 shed virus at 3 dpi and peaked at 5 dpi. Thus, we speculate that the fast replication of IAV in pigs may have rapidly stimulated proinflammatory responses in the upper respiratory tracts and consequently inhibited the replication of IDV in pigs. Of interest, interference of IAV on IDV was not observed in the tissues of the middle or lower respiratory tracts, perhaps due to slower proinflammatory responses of IAVs within the lower respiratory tract, compared with those at the upper respiratory tracts. Future studies will include the collection of swine bronchoalveolar lavage fluid to evaluate the proinflammatory responses during different days of coinfection. Nevertheless, our data support that virus-virus interactions during coinfection are complicated and likely affected by multiple factors such as the time lag between coinfecting viruses and rate of virus replication (reviewed in (34)).

Virus-virus interaction, via incredibly diverse mechanisms, are broadly classified by three outcomes: interference, enhancement, or accommodation (34, 35), and these interactions can be groups into 15 mechanisms with three main categories, direct interactions between the viruses, indirect interactions that result from alterations in the host environment, and immunological interactions (reviewed by DePalma *et al.* (35)]. The most frequently observed interaction is interference, or when replication of one virus prevents or inhibits multiplication of the other. Viral interference has also been defined as a state of temporary immunity from infection induced by viral infection (36), and the most common mechanism of viral interference is interferon mediated. One virus triggers the host interferon response that nonspecifically blocks replication of the other virus. Time of exposure and viral replication are critical factors. On the other hand, viruses may also compete for receptor binding or replication sites, metabolites, or other host supports, and this competition can occur between closely related or unrelated viruses. Our results suggested that the interactions between IAV and IDV were associated with the proinflammatory responses by either virus, or the inhibitory interference were shown to be bi-directional. Our study also showed that IAV and IDV did not interfere each other if both viruses were inoculated within a certain time frame, e.g., 24 hours when IAV inoculation followed by IDV inoculation or 48 hours when IDV inoculation followed by IAV inoculation. As described before, such time gaps are more likely associated with the speed of virus replication as well as the proinflammatory responses induced by the first virus.

Several prior studies detail IAV stimulated proinflammatory responses in humans, various animal models and in various cells (29). For example, in newborn pig trachea cells, subtype H3N2 swine IAV activated JAK/STAT and MAPK signaling pathways and stimulated the upregulation of RIG-I, IFN-β, IFN-Λ1, Mx1, OAS1, PKR, IL6, and SOCS1 (37). On the other hand, lung tissues from the mice infected with IDV had minor proinflammatory responses for TLR7, CCL5, IRF3, IL-6, IL-1β, IFN-γ at 1 dpi (27), of which, CCL5 had the highest responses. Of interest, although IAV can lead much higher morbidity in pigs than IDV, our *in vitro* study using STEC showed both viruses can induce a similar level of proinflammatory responses with the same set of genes, including IP-10, CCL5, CXCL9, TNF-α, and IFN-β. Several markers evaluated, IL-4, IL-6, IL-8, IL-10, and IL-1β, had minimal or limited expression. These results indicate IDV and IAV may share similar signaling pathways, such as JAK/STAT and MAPK, during proinflammatory immune responses.

Apoptosis, or programmed cell death in the absence of inflammation, is an energy-dependent, caspase-mediated biochemical mechanism characterized morphologically by cytoplasmic and nuclear condensation, chromatin cleavage, apoptotic bodies, maintenance of an intact plasma membrane, and exposure of surface molecules targeting phagocytosis and efficient removal of the cell and its contents (38–41). Activation of CC3 is shown to be essential for efficient influenza virus propagation (33). The induction of apoptosis and subsequent phagocytosis of infected cells is also one of antiviral mechanisms (42, 43). In this study, we evaluated the CC3 expression in trachea which showed that the average number of CC3 positive cells were higher in the trachea from the pigs from all pig treatment groups than those from the negative control pig, indicating virus caused apoptosis in the infected pigs (Fig. 5E). On the other hand, no statistical significance was found among A-mock, D-mock and A+D groups. The results support that coinfection pigs did not have increased pathogenesis than either IAV or IDV single infection pigs.

One limitation of this study is that only a single dose for each virus was used both *in vitro* and *in vivo* experiments. Additional experiments need to further evaluate alternative doses for inoculation, which may affect the interference patterns between two viruses. In addition, in the pig experiment, we only performed simultaneous coinfection in the pig model, and the primary proinflammatory cytokines were not determined. Future directions could feasibly test a sequential infection time course in the pig model, and the tissue dependent primary proinflammatory cytokines aid in further understanding of the tissue dependent virus interference in pigs.

In summary, this study suggests that both IAV and IDV can interfere with the replication of each other by stimulating proinflammatory responses; however, the proinflammatory response was 24 hours slower for IDV than IAV. The mechanism of viral interference appears to be via proinflammatory responses and not through viral binding or replication. Coinfection of IDV and IAV in pigs did not show enhanced pathogenesis, compared with those infected only with IAV. This study facilitates our understanding of virus epidemiology and pathogenesis associated with IAV and IDV coinfection.

## Materials and methods

### Viruses and Cells

Animals were infected with D/bovine/C00046N/Mississippi/2014 (abbreviated as D/46N) and/or A/swine/Texas/A01104013/2012 (H3N2) (abbreviated as sH3N2) isolated from feral swine. D/46N was isolated from sick cattle in Mississippi (44) and propagated in human rectal tumor cells (HRT-18G) (American Type Culture Collection, Manassas, VA), whereas sH3N2 was isolated from feral swine (45) and propagated in MDCK cells (American Type Culture Collection, Manassas, VA). Viruses were propagated in Opti-MEM I Reduced Serum Medium (Thermo Fisher Scientific, Asheville, NC) supplemented with 1 μg/ml of TPCK-trypsin (Gibco, New York) at 37°C under 5% CO_2_. The predominant exposure for swine in the U.S. is the IAV H3 subtype and strains circulating in feral and domestic swine are antigenically and genetically similar (45–48). Because pathogenicity is strain, dose and route dependent, we used swine IAV and IDV stains previously shown by our laboratory to produce successful infection in pigs at 10^6^ TCID_50_/ml by intranasal inoculation (25, 49).

### Growth kinetics *in vitro*

To determine the replication consequences of the coinfection in cell lines, STEC in 6-well plates were infected with viruses at a multiplicity of infection of 0.001 for IAV and 0.1 for IDV, respectively, based on our pilot study. For the four groups of single infection, after absorption for 1 hour at 37°C, the cells were washed with PBS and incubated for 96 hours at 37°C in 5% CO_2_ with Opti-MEM I Reduced Serum Medium (Thermo Fisher Scientific, Waltham, MA) supplemented with 1 μg/ml TPCK treated Trypsin from bovine pancreas (Sigma-Aldrich, St. Louis, MO) and with 100 U/ml Gibco penicillin-streptomycin (Thermo Fisher Scientific, Waltham, MA). For the multiple-infection groups, supernatants were removed, washed with PBS, and re-infected again at designated hours (6h, 24h and 48h) after first infection. Supernatants were collected at 24, 48, 72 and 96 hours after infection. The RNA copies of each samples were determined with qRT-PCR. STEC of 24-, 48- and 72-hours post infection were washed, harvested and subjected to total RNA extraction and mRNA expression analyses of cytokines and chemokines.

### Quantification of cytokine and chemokine expression

RNeasy Mini Kit (QIAGEN, Germantown, MD) was used to extract total RNA from infected cells following the manufacturing manual. Total cellular RNA of 1μg was transcribed to cDNA using SuperScript™ III Reverse Transcriptase (Thermo Fisher Scientific, Waltham, MA) with Oligo(dT)20 Primer (Thermo Fisher Scientific, Waltham, MA, USA). The cDNA was used in qPCR using PowerUp™ SYBR® Green Master Mix (Thermo Fisher Scientific, Waltham, MA) and designed primers for specific targets (Table 1). The qPCR amplification mixture contains: 7μl of water, 10μl of PowerUp™ SYBR® Green Master Mix, 1μl of each forward- and reverse-primers (10 μM), 1μl of cDNA. The parameters of the qPCR were as follows: one cycle at 50°C for 2 minutes, one cycle at 95°C for 2 minutes, followed by 40 cycles at 95°C for 1 seconds, 60°C for 30 seconds. Gene expression data were normalized by house-keeping gene (β-actin). We used the 2^-ΔΔCt^ (Ct is the cycle threshold) methods for qPCR data analysis. Here, ΔΔCt represents: ΔCt (sample) ([Ct_gene of interest_^-Ct^_housekeeping gene_] of infected cells) - ΔCt (Mock) ([Ct_gene of interest_ -Ct _housekeeping gene_] of uninfected cells). The mean fold change (2^-ΔΔCt^) values of triplicates and standard deviation were represented. Porcine IFN-β ELISA Kit (Abcam, Cambridge, MA) was used to quantify the protein expression of IFN-β in supernatants from STEC infection following the manufacturing manual.

### Viral RNA extraction

The supernatant from homogenized tissue and the transport media containing the nasal swabs were used for RNA extraction. Viral RNA was extracted using the MagMAX Pathogen RNA/DNA Kit (# 4462359) with KingFisher™ Flex Purification System (Thermo Fisher Scientific, Waltham, MA) following the manufacturer’s high-throughput purification protocol. Extracted RNA was stored at −80°C until qRT-PCR could be performed.

### Virus quantification

To quantify viral copy number in nasal swabs and tissues, qRT-PCR was performed using standard protocols, primers, and probe validated by the CDC for IAV (50) and in-house designed primers and probe to detect IDV (below). Briefly, qRT-PCR was performed in triplicate by using TaqMan Fast Virus 1-step Master Mix (Life Technology, Carlsbad, CA) following the manufacturer’s protocol and using 2 μl of RNA template. Samples were amplified using IAV CDC primer and probe set InfA: Forward 5’-GACCRATCCTGTCACCTCTGAC-3’; Reverse 5’-AGGGCATTYTGGACAAA CGTCTA-3’; and Probe 5’-[FAM]-TGCAGTCCTCGCTCA CTGGGCACG-[BHQ]-3’. To detect IDV primer and probe set Forward 5’-ACGCAATGGCACAAGAAC-3’; Reverse 5’-ACCACTATGCTCTCTCCAC-3’; and Probe 5’-[FAM]-AGGAGTTAACCCAATGACCAGGCAAACGA-[BHQ]-3’ was used. The fast mode amplification protocol was followed: reverse transcription (1 cycle at 50°C for 5 min), inactivation (1 cycle at 95°C for 20 sec), followed by 40 alternating cycles of denaturation at 95°C for 3 sec and annealing and extension at 60°C for 1 min.

Viral copies in samples were determined with the standard curve generated by the plasmid containing the target gene segment (IAV M plasmid or IDV M plasmid) cloned into a dual-promoter plasmid vector, pHW2000, as previously described (51, 52). The IDV M plasmid was generously provided by Dr. Richard Webby (St. Jude Children’s Research Hospital, Memphis, TN). The standard curves were plotted by Ct values against viral copy number/ml (nasal swabs) or viral copy number/g (tissue homogenate). Mean Ct values of biological triplicates were recorded, and viral copy number concentrations were calculated based on the standard curve constructed across a series of known target concentrations of plasmid. The data were presented in figures as log 10 (viral copy number concentration) form.

### Animal study

Animal experiments were conducted under BSL-2 conditions in compliance with protocols approved by the Institutional Animal Care and Use Committee of Mississippi State University. Twenty-five pigs (Large White x Landrace) aged 116-120 days with a mean weight of 43 kg (range 24-62 kg) were provided by the Mississippi State University Department of Animal and Dairy Sciences (MSU-ADS). The pigs farrowed at MSU-ADS were unvaccinated and biosecurity protocols were in place to limit contact with other animals and personnel with self-reported clinical symptoms of respiratory infection as well other animals. All pigs were housed together prior to the study. At - 8 dpi and repeated at 0 dpi, all pigs were confirmed serologically negative by HI assay against the challenge viruses and other representative influenza viruses. These viruses were chosen to represent the antigenic strains in the vaccine used to vaccinate the sows (Flusure XP® Zoetis, USA) at 84 days gestation as well as seasonal human influenza strains. The pigs tested seronegative against the following viruses: D/46N (challenge virus), sH3N2 (challenge virus), A/swine/Ohio/09SW96/2009 (H3N2), A/swine/Indiana/13TOSU1154/2013 (H1N1), A/swine/Iowa/15/2013 (H1N1), and human viruses A/Hong Kong/4801/2014 (H3N2) and A/California/04/2009 (H1N1).

At 5 days prior to inoculation all pigs were transferred to BSL-2 facilities and assigned by ear tag number to one of four treatment rooms (12 x 12 feet with negative air flow) as indicated below. Although all pigs were 116-120 days old, their weight range was variable, so they were first stratified in groups of four of similar weights and then randomly assigned to one of four infection treatment groups: A-mock group (sH3N2) (n=5), D-mock group (D/46N) (n=7), IAV + IDV group (coinfection) (n=7) or negative group (sterile PBS) (n=6). Investigators and animal care personnel were not blinded to treatment groups and workflow was control, IDV, IAV and coinfection Pigs were intranasally infected as follows: A-mock group received 10^6^ TCID_50_/ml of sH3N2 in a volume of 1 ml administered in approximately equal doses to the right and left nostril by syringe; D-mock group received 10^6^ TCID_50_/ml of D/46N using the same method as above; A+D group received 10^6^ TCID_50_/ml of sH3N2 and 10^6^ TCID_50_/ml of D/46N using the same method as above; and negative group received sterile PBS using the same method as above. Because pathogenicity is strain, dose and route dependent, we used swine IAV and IDV stains previously shown by our laboratory to produce successful infection in pigs at a smaller dose (10^6^ TCID50/ml) and by intranasal inoculation (25, 49). IAV H3 subtype is the predominant influenza virus exposure in feral and domestic swine (46–48) and intranasal inoculation simulates a more natural route of infection (53).

During the study, clinical signs, rectal temperatures, and nasal swabs were taken daily and whole blood collected at 0, 3 and 5 dpi. At 5 dpi all pigs were euthanized, nasal swabs and blood were collected immediately prior to the euthanasia. Pigs were necropsied and respiratory tract tissues were collected including: rostral, middle and ethmoid sections of nasal turbinate; soft palate; upper, middle and distal sections of trachea; bronchus; and one section from each lung lobe (left cranial, left caudal, right cranial, right middle, right accessory and right caudal). Tissues were fixed in 10% buffered formalin and additional sets were frozen at −80°C.

### Clinical Data

To assess clinical signs of influenza infection, prior to entering the enclosure, pigs were observed from a window for changes in attitude, elevated respiratory rate, cough, dyspnea, nasal or ocular discharge or conjunctivitis. Rectal temperatures were obtained for all pigs beginning three days prior to inoculation (−3 dpi) and daily through day 5 of the study (5 dpi). Nasal swabs were collected daily (0-5 dpi) using sterile cotton tipped applicators and transported in sterile PBS supplemented with PenStrep (1:100 w/v) on ice to the BSL-2 laboratory where they were aliquoted and stored at −80°C. Blood samples were taken at 0, 3 and 5 dpi and stored at 4°C until a complete blood count (CBC) could be performed by the MSU-CVM Diagnostic Lab. A CBC for each pig was obtained with exception of blood samples that were clotted prior to processing. The samples included one pig from the negative group at 0 dpi, one pig from the D-mock group at 0 dpi, one pig from A+D group at 3 dpi and at 5dpi and one pig from the A-mock group at 5dpi.

Respiratory tract tissues were collected at necropsy and frozen at −80°C until homogenization. Tissues were thawed on ice and a sterile #10 blade and forceps were used to cut and then weigh 1 gram of tissue. Tissue samples were placed into prechilled 7 ml autoclaved tubes with prefilled ceramic beads (KT03961-1-302.7, Bertin Instruments, Rockville, MD) and 4 ml of prechilled PBS supplemented with PenStrep (1:100 w/v). Tissues were homogenized at 8000 x rpm for 20 seconds for 4 cycles (Precellys® Evolution Homogenizer, Bertin Instruments, Rockville, MD). Sample heating was prevented by incubating the tubes on ice between homogenization cycles. Samples were centrifuged at 15871 x g (Eppendorf ® 5424, Eppendorf North America, Hauppauge, NY) for 5 minutes to pellet debris and the supernatant aliquots stored at −80°C until RNA extraction and qRT-PCR could be performed.

### Histopathological examination

Respiratory tract tissues were fixed in 10% buffered formalin, paraffin embedded, sectioned at 5μm sections and stained with hematoxylin and eosin (H&E) for histopathological examination.

### Caspase-3 stain and quantification

Tracheal sections cut at 5μm on charged slides. Slides were stained with anti-cleaved caspase-3 (Asp175) antibody (Cell Signaling Technology, Catalog #9661) at a 1:200 dilution following the manufacturer’s protocol for IHC paraffin-embedded tissues. The tissue sections were evaluated at 20X to determine the area with the most abundant staining, and then positive cells from 20 consecutive high powered fields (40X) were counted. The average number of CC3 positive staining cells from the upper, middle and distal trachea were recorded and analyzed.

### Statistical analyses

One-way analysis of variance (ANOVA) with repeated measures was used to compare the growth kinetics in cells, with Bonferroni adjustment for multiple comparisons (GraphPad Prism version 8.3.1, GraphPad Software, San Diego, CA). Two-way ANOVA was conducted for the animal study with a replication comparison between the single infection groups and co-infection group followed by Bonferroni multiple comparisons (GraphPad Prism version 8.3.1, GraphPad Software, San Diego, CA). The CC3 data were log10-transformed and subjected to the Shapiro–Wilk’s test of normality and Brown-Forsythe test for homogeneity of variance. A One-way ANOVA followed by Tukey’s multiple-comparisons test was performed. Differences were considered significant when *p* ≤ 0.05.

## Acknowledgments

We thank Dr. Stacey Schultz-Cherry for kindly providing swine tracheal epithelial primary cells (STEC) and Kaitlyn Waters and Hui Wang for their help with the animal experiments. We also thank the Mississippi State University College of Veterinary Medicine’s laboratory animal resources and care to support our animal experiments.

